# Protocatechuic acid induces endogenous oxidative stress in CR-hvKP by regulating the EMP-PPP pathway

**DOI:** 10.1101/2024.03.06.583678

**Authors:** Yesheng Zhong, Yumeng Cheng, Shuai Xing, Xiaoxiao Zhang, Shiqi Luo, Xinru Shi, Yang He, Huixin Liu, Meng Yang, Hongbin Si

**Affiliations:** College of AnimaScience and Technology, Guangxi KeyLaboratory of Animal Breeding, DiseaseControl and Prevention, Nanning, 530004, PRChina; College of Animal Sciences, Fujian Agriculture and Forestry University, Fuzhou, 350002, China; College of Life Sciences, Northwest A&F University, Yangling, Shaanxi, 712100, China

**Author notes:** Corresponding Author: Prof: Hongbin Si.

**Keywords:** Carbapenem-resistant hypervirulent Klebsiella pneumoniae, Protocatechuic acid, Glycolysis, Pentose phosphate pathway, Oxidative stress

## Abstract

**Background:** Klebsiella pneumoniae is an important opportunistic pathogen and zoonotic pathogen. The widespread use of antibiotics has led to the emergence of a large number of multidrug-resistant Klebsiella pneumoniae in clinical animal husbandry, posing a serious threat to global health security. Protocatechuic acid (PCA) is a phenolic acid substance naturally present in many vegetables and fruits. It is a safe and highly developed new type of antibacterial synergist.

**Purpose:** This study explored the antibacterial and synergistic mechanisms of PCA against Carbapenem-resistant hypervirulent Klebsiella pneumoniae.

**Study design:** Metabolomic analysis using PCA to investigate the metabolic effects of CR-hvKP and further explore the antibacterial mechanisms resulting from this metabolic regulation.

**Methods:** The MIC of PCA was measured by microdilution, and its bactericidal effect was observed by DAPI staining. Resistance and hemolysis tests were performed to ensure safety. The synergy of PCA and meropenem was tested by checkerboard assay. The biofilm inhibition was assessed by crystal violet and EPS assays. The membrane morphology, permeability, and potential were examined by SEM, PI, NPN, and DiSC3(5). The metabolic changes were evaluated by AlamarBlue, metabolomics, enzyme activity, ELISA, molecular docking, and qRT-PCR. The oxidative stress and metabolic disorders were verified by NADP(H), ROS, MDA, and ATP assays.

**Results:** The results showed that PCA can synergize with antibiotics and inhibit the biofilm and membrane functions of CR-hvKP at low concentrations. Metabolomics revealed that PCA affects the EMP and PPP pathways of CR-hvKP, causing oxidative stress. This involves the binding of PGAM and the downregulation of BPGM, leading to the accumulation of glycerate-3P. This results in the inhibition of G6PDH and the imbalance of NADPH/NADP+, disrupting the energy metabolism and increasing the oxidative stress, which impair the biofilm and membrane functions and enhance the antibiotic efficacy.

**Conclusion:** The results demonstrate that PCA regulates the EMP-linked PPP pathway of CR-hvKP, inhibits biofilm and membrane functions, and synergizes with antibiotics to kill bacteria, providing new insights and candidates for natural antibacterial enhancers.

**Author Summary:** Klebsiella pneumoniae is a common pathogenic bacterium that can infect both humans and animals, causing serious diseases such as pneumonia, meningitis, and sepsis. Due to the overuse of antibiotics, this bacterium has developed resistance to many drugs, posing a significant threat to global health security. Through our research, we have discovered a natural substance called protocatechuic acid (PCA) that can enhance the effectiveness of antibiotics against this bacterium. PCA is found in many vegetables and fruits and is a safe and non-toxic antibacterial adjuvant. Our analysis of the metabolomics of PCA on Klebsiella pneumoniae has revealed its antibacterial and synergistic mechanisms. The study found that PCA can affect the bacterium’s sugar metabolism pathway, leading to the generation of endogenous oxidative stress. This disrupts their energy metabolism, damages their cell membranes and biofilms, making them more susceptible to being killed by antibiotics. Through this mechanism, PCA can synergize with common antibiotics such as meropenem, enhancing their bactericidal ability. Our research has demonstrated that PCA is an effective antibacterial adjuvant, providing new candidates and insights for the development of natural antibacterial agents.

**Graphical abstract:** 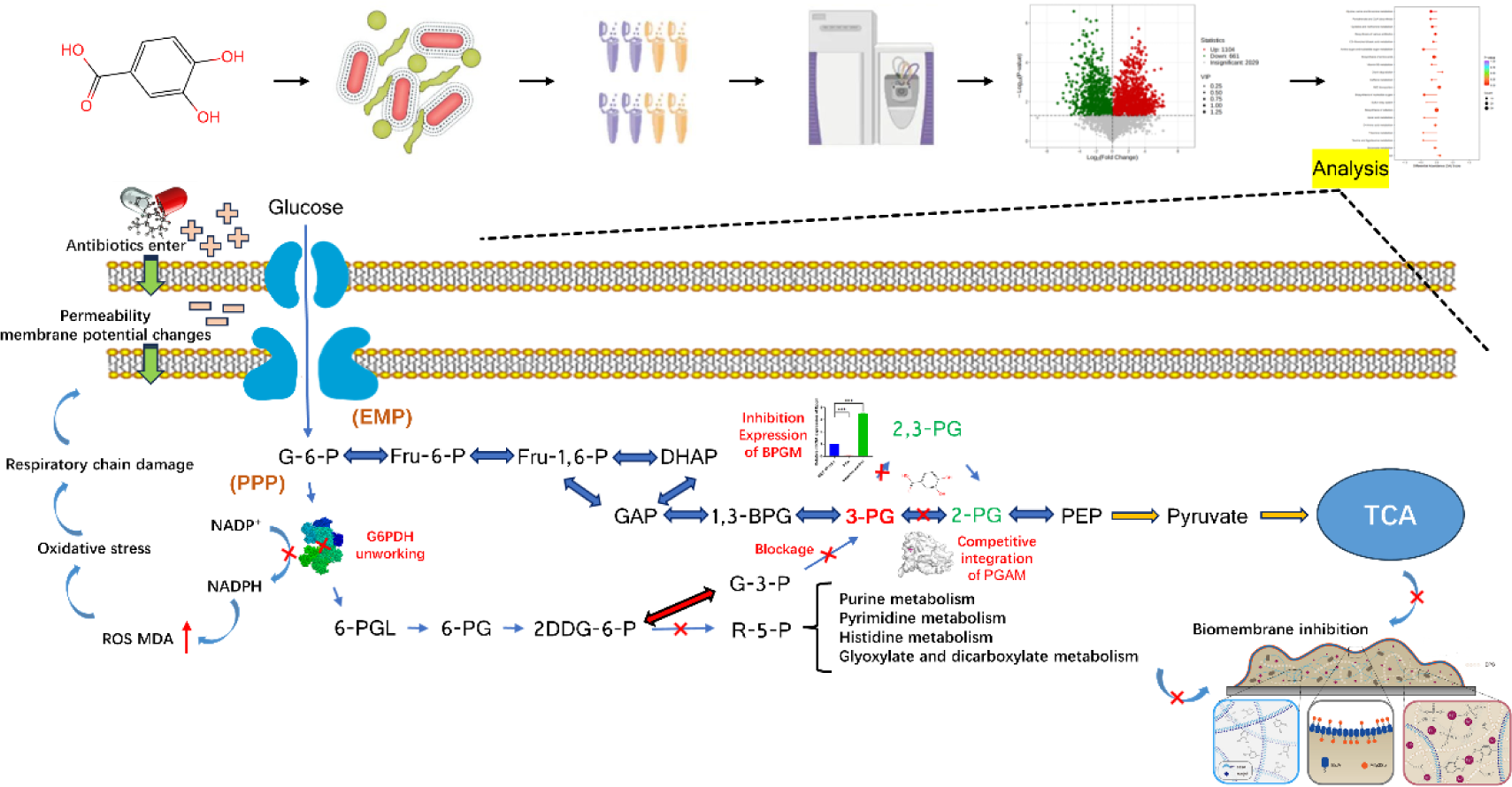

## 1 Introduction

Klebsiella pneumoniae is a Gram-negative bacterium widely distributed in the skin, respiratory and digestive tracts, urinary and reproductive systems of humans and animals, as well as in the natural environment. It is an important opportunistic pathogen and zoonotic pathogen, capable of causing severe infectious diseases such as pneumonia, sepsis, urinary tract infections, and meningitis. It can be transmitted to humans and animals through the food chain or direct contact, posing a threat to public health and animal husbandry[1]. Due to the widespread use and abuse of broad-spectrum antibiotics, Klebsiella pneumoniae has developed multiple drug resistance, especially resistance to carbapenems, making the treatment of Klebsiella pneumoniae infections extremely difficult. In recent years, genetic elements involved in the exchange of carbapenemase-producing cKP and hvKP have been discovered, indicating the emergence of carbapenem-resistant hypervirulent Klebsiella pneumoniae (CR-hvKP). CR-hvKP not only produces carbapenemases, leading to resistance to carbapenems, but also expresses various virulence factors such as collagenase, siderophore, and capsule, enhancing its invasiveness and lethality [3–4]. CR-hvKP has been reported in multiple countries and regions worldwide, becoming a new clinical challenge and public health threat [4].

Therefore, it is urgently needed to search for new antibacterial agents to combat CR-hvKP infection. Natural plant antibacterial agents refer to substances with antibacterial activity extracted or synthesized from plants, such as flavonoids, phenolic acids, tannins, anthocyanins, etc. These substances have the characteristics of being natural, safe, non-toxic, and free of side effects, and will not cause harm to the human body and animals [5]. Natural plant antibacterial agents have multi-target and multi-mechanism antibacterial properties, which can affect the growth and reproduction of bacteria in different ways, such as destroying cell walls and membranes, changing membrane potential and energy metabolism, and inhibiting the synthesis of biomacromolecules [6]. These methods can effectively combat infections caused by multidrug-resistant bacteria and provide new strategies for the treatment of refractory diseases. Protocatechuic acid (PCA), also known as 3,4-dihydroxybenzoic acid, is a phenolic acid substance naturally present in many vegetables and fruits, such as blackberries, strawberries, cherries, grapes, tea, etc. [7]. PCA has various biological activities such as antioxidant, anti-inflammatory, and anti-tumor effects, which can eliminate free radicals, inhibit the release of inflammatory factors, and induce apoptosis of cancer cells [8]. PCA also has certain antibacterial effects and can inhibit the growth of various bacteria, such as Staphylococcus aureus, Escherichia coli, Salmonella, etc. [9–10]. Moreover, PCA can inhibit the biofilm formation of Klebsiella pneumoniae by inhibiting its quorum sensing [11]. However, the antibacterial effect of PCA on CR-hvKP and its mechanism have not been fully studied.

In this study, we demonstrated that PCA promotes the disruption of intracellular redox balance by affecting key enzymes in the CR-hvKP metabolic process, leading to changes in membrane permeability and inhibition of biofilm function, ultimately synergizing with antibiotics to kill drug-resistant bacteria.

## 2 Materials and Methods

### 2.1 Bacteria strains

Klebsiella pneumoniae (MEC19116-1) and Klebsiella pneumoniae (FMEC20113-2), these two strains are carbapenem-resistant Klebsiella pneumoniae (CRKP) selected from samples of edible animals and their surrounding growth environment. After preliminary identification of virulence genes, both strains were found to have highly virulent genes. They were cultured in LB medium at 37°C.

### 2.2 Chemicals and reagents

Protocatechuic acid is purchased from Hengrui Tongda Company. Congo red, sucrose, glucose, and KCl are products of Glucose Company. Dimethyl sulfoxide, meropenem, tigecycline, dexamethasone, DAPI, PI, HEPES buffer, AlamarBlue, ROS detection kit, NADP(H) detection kit, and G6PDH enzyme detection kit are products of Solarbio Company. ATP detection kit and MDA detection kit are products of Biyun Tian Company. DisC3(5) and NPN are products of Aladdin Company. Glycerate-3P is a product of Shanghai Yuan Ye Biotechnology Co., Ltd. PGAM enzyme-linked immunosorbent assay kit and BPGM enzyme-linked immunosorbent assay kit are products of MEIMIAN Company. Bacteria RNA Extraction Kit, reverse transcription kit, and qRT-PCR detection dye are products of Genstar Company.

### 2.3 Minimum inhibitory concentration and minimum bactericidal concentration

Add an appropriate concentration of PCA to the liquid culture medium and dilute it into different concentrations of drug culture medium. Take a certain volume of bacterial suspension and mix it with each concentration of medium in a 96-well plate. Place the prepared 96-well plate in a bacterial incubator for overnight incubation. Based on the growth of bacteria in the 96-well plate, the drug concentration corresponding to the last area without white spots of bacterial growth can determine the minimum inhibitory concentration (MIC) of the drug. To determine the minimum bactericidal concentration (MBC), take 10μL of the culture fluid from the MIC well and the two wells on the left side of the drug concentration, and inoculate them on a drug-free agar plate. After average spreading, incubate again overnight in a 35℃ incubator with normal air. The lowest drug concentration required to kill 99.9% (reducing by 3 orders of magnitude) of the test microorganisms is defined as the MBC [12].

### 2.4 Time-dependent inhibition curve

Prepare LB medium according to the preparation method and sterilize it. Add different concentrations of PCA and inoculate bacteria in the logarithmic growth phase into the medium. Place the inoculated medium in a constant temperature shaker for cultivation. Use a spectrophotometer to regularly measure the bacterial quantity. Based on the measured bacterial quantity data, plot the bacterial growth curve [13].

### 2.5 DAPI staining

After culturing bacteria to the logarithmic growth phase, different concentrations of PCA were added and incubated for 30 minutes. The cells were then collected by centrifugation at 4℃ and 5000rpm for 5 minutes. The cells were washed three times with PBS buffer and DAPI was added to precipitate. Subsequently, the cells were treated in the dark for 20 minutes, followed by centrifugation at 5000rpm for 3 minutes and three additional washes with PBS buffer. The cells were resuspended in 100μL of PBS buffer and 10μL was taken onto a glass slide. A cover glass was placed on top and the cells were observed under a fluorescence microscope and photographed [14].

### 2.6 Hemolysis assay

Take 5mL of mammalian blood using an anticoagulant tube, centrifuge at 1000g for 10 minutes under a frozen centrifuge, pour off the supernatant, slowly add physiological saline and invert repeatedly until well mixed, repeat centrifugation 2-3 times until a clear supernatant is obtained, and dilute it to a 5% red blood cell suspension. Dilute the original chlorogenic acid to different concentrations of drug groups using physiological saline, and set up a physiological saline group and a prednisolone group. Mix the red blood cell suspension with the drug solution and add it to a 96-well cell plate, incubate at 37°C on a shaker for 1 hour, centrifuge at 1000g for 10 minutes at 4°C, transfer a certain amount of supernatant to another 96-well cell plate, and add an equal volume of physiological saline. Measure the absorbance at 540nm [15].

### 2.7 Research on the development of drug resistance

The overnight shaken bacteria were inoculated into a new broth culture medium and added with 1/2 MIC concentration of PCA and positive control drug tetracycline, and incubated on a shaker for 12 hours as one generation. Then, the bacteria were taken out for drug sensitivity test and continued to be subcultured. The concentration of the drugs added should be maintained at ½ MIC. Finally, it was subcultured until the twentieth generation and a drug sensitivity curve was plotted [16].

### 2.8 Checkerboard experiments

The PCA and meropenem were separately diluted in a 96-well plate, and then they were mixed in equal proportions to form multiple drug combinations with different concentrations. An equal volume of bacterial suspension was added afterwards. Positive and negative control groups need to be designed in the experiment. The prepared 96-well plate was placed in a bacterial incubator for overnight incubation, and the drug concentration corresponding to the area without white spots in the 96-well plate was recorded. The formula for the result is FICI = MICab/MICa + MICba/MICb [17].

### 2.9 Characterization of metabolism observations

Using 0.08g of Congo red, 5g of sucrose, 1g of agar, 5g of brain-heart infusion medium, 200mL of deionized water, and different concentrations of PCA as Congo red medium, the bacteria were inoculated in liquid medium containing 5% glucose and cultured for 24 hours. Then, 10μL of bacterial suspension was taken and inoculated onto Congo red solid medium, followed by incubation at 37℃ for 24 hours. Afterwards, the plates were taken out and observed at room temperature. If the colonies appeared black, it indicated that the bacteria were biofilm-positive; if the colonies appeared red, it indicated that the bacteria were biofilm-negative [18].

### 2.10 Crystal violet staining

Bacteria with a logarithmic growth phase were inoculated into a 96-well plate containing LB medium at a concentration of 10^6 CFU/mL, with 200μL per well. The plate was set up with a positive control group (containing only bacteria) and different concentrations of PCA groups. The 96-well plate was incubated in a constant temperature incubator at 37°C for 48 hours to allow the bacteria to form a biofilm at the bottom of the wells. The plate was then removed from the incubator, gently tilted, and the culture medium was discarded by shaking. Any remaining liquid was removed using a sterile pipette. Each well was washed three times with sterile PBS buffer, with 200μL per wash, to remove non-adherent cells and impurities. 200μL of a 1% crystal violet solution was added to each well and stained at room temperature for 15 minutes. The crystal violet staining solution was then poured out, and each well was rinsed with tap water until no color overflowed, to remove excess dye. The 96-well plate was inverted and dried on absorbent paper at room temperature or dried at 37°C. 200μL of a 33% acetic acid solution was added to each well and allowed to act at room temperature for 15 minutes to dissolve the crystal violet. The absorbance of each well was measured using an enzyme-linked immunosorbent assay (ELISA) reader at a wavelength of 570nm [19].

### 2.11 Extracellular polymer content determination test

Cultivate bacteria to the logarithmic growth phase, dilute 100 times and take 2mL of bacteria into a 6-well plate. Place the cell crawler into the well and introduce varying concentrations of PCA. Culture for 48 hours, then use tweezers to remove the scraper and transfer it to a 50mL Eppendorf tube. Perform ice bath sonication for 30 minutes at 4°C, then centrifuge at 4000rpm for 30 minutes under the same conditions. Transfer the supernatant to a 2mL glass tube, add 1mL of phenol and 5mL of concentrated sulfuric acid, and react in the dark for 30 minutes. Measure the absorbance at 490nm wavelength [20].

### 2.12 Scanning Electron Microscope

Bacteria collected by centrifugation at 3500rpm for 10 minutes and cultured to logarithmic growth phase were resuspended in 1×PBS (PH7.4) three times to OD600=0.5. Then, different concentrations of PCA were added for experimental grouping as shown in the table below, and each test tube was incubated at 37℃ for 4 hours. After incubation, the bacterial suspension was centrifuged at 4000rpm for 10 minutes, resuspended in 1×PBS (PH7.4) three times, and the supernatant was removed to collect the precipitate. The precipitate was fixed in 2.5% glutaraldehyde solution for 2 hours, then washed with PBS to remove the fixative, and sequentially placed in 40%, 50%, 60%, 70%, 80%, and 90% ethanol for 15 minutes for gradient dehydration. It was then replaced with 100% tert-butanol for 30 minutes, dried, gold-coated, and observed and photographed using a scanning electron microscope [21].

### 2.13 Inner membrane permeability

After culturing bacteria to the logarithmic growth phase, different concentrations of PCA were added and incubated for 30 minutes. Then, 10μL of PI staining solution was added and incubated for another 30 minutes under dark conditions. The samples were transferred to a 96-well cell plate and the fluorescence intensity was measured using an enzyme labeling instrument (excitation wavelength: 535nm; emission wavelength: 620nm) [22].

### 2.14 Outermembranepermeability

Bacteria in the logarithmic growth phase were collected by low-speed centrifugation, washed twice with HEPES buffer (5mM HEPES + 5mM Glucose), resuspended, and then different concentrations of PCA were added. A negative control group was also set up. After a 30-minute treatment, 90μL of the sample was taken and added to a 96-well cell plate, followed by the addition of 10μL of NPN for mixing. The fluorescence intensity was measured using an enzyme labeling instrument (excitation wavelength: 350nm; emission wavelength: 420nm) [22].

### 2.15 Membrane depolarization assay

Bacteria in the logarithmic growth phase were collected by low-speed centrifugation, washed twice with HEPES buffer (5mM HEPES + 20mM Glucose), and then resuspended. Subsequently, 10μL of KCL and 10μL of DisC3(5) were added to each sample tube, followed by gentle mixing after a few minutes. Different concentrations of PCA were added to each sample tube, with a negative control group set. The samples were treated for 30 minutes, and then transferred to a 96-well cell plate. Fluorescence intensity was measured using an enzyme labeling instrument (excitation wavelength: 622nm; emission wavelength: 670nm) [23].

### 2.16 Bacterial metabolic viability experiment

Bacteria with a logarithmic growth phase were inoculated into a 96-well plate containing LB medium at a concentration of 10^6 CFU/mL, with 200μL per well. Blank control group (containing only LB medium) and positive control group (without drugs) were set up, and different concentrations of PCA were added. The 96-well plate was incubated in a constant temperature incubator at 37°C for 1 hour to allow the bacteria to adapt to the environment. 10μL of alamarBlue reagent was added to each well to achieve a final concentration of 10%. The 96-well plate was incubated in a constant temperature incubator at 37°C for 4 hours to allow the alamarBlue reagent to react with the bacteria. The absorbance (at a wavelength of 570nm) and fluorescence intensity (excitation wavelength of 530nm, emission wavelength of 590nm) of each well were measured using an enzyme-linked immunosorbent assay reader, and the relative absorbance (subtracting the absorbance of the blank control group) and relative fluorescence intensity (subtracting the fluorescence intensity of the blank control group) were calculated. The metabolic activity of bacteria at different drug concentrations was evaluated using relative absorbance or relative fluorescence intensity as indicators and compared with the positive control group [24].

### 2.17 Untargeted metabolomics

According to the preliminary experimental results, the concentration of PCA was selected to study the sub-inhibitory effect on CR-hvKP, and metabolomics analysis was conducted. The original catechin with or without sub-inhibitory concentration was added to the logarithmic growth phase CR-hvKP bacterial suspension and incubated at 37°C with constant shaking for 30 minutes. The bacteria were then collected by centrifugation for 10 minutes, washed with sterile PBS, and stored in liquid nitrogen. Metabolite samples were extracted from the intracellular and extracellular samples using methanol/water (8:2, v/v). The intracellular and extracellular metabolite samples were analyzed using liquid chromatography-mass spectrometry (LC-MS) and gas chromatography-mass spectrometry (GC-MS), respectively. Data acquisition and processing were performed using Xcalibur software, and data analysis and bioinformatics annotation were conducted using MetaboAnalyst software [25].

### 2.18 G6PDH enzyme activity assay

Bacteria were cultured to the logarithmic growth phase and divided into two groups. One group was treated with different concentrations of PCA, while the other group was treated with different concentrations of glycerate-3P. After a 30-minute treatment, 1mL of extraction solution was added, followed by ultrasonic disruption of the bacteria (200w, 3s ultrasound, 10s interval, repeated 30 times). The mixture was then centrifuged at 8000g for 10 minutes at 4℃, and the supernatant was collected as the sample. In blank tubes and test tubes, 50μL of distilled water and 50μL of the sample were added respectively, along with 950μL of working solution. 200μL of the mixture was transferred to a 96-well plate, and the absorbance at 340nm was measured for both groups within 5 minutes. G6PDH activity (U/104cell) = 1.286×A2(300s)-A1(0s)[26].

### 2.19 Enzyme-linked immunosorbent assay

Dilute the capture antibody (1:100) in coating buffer (10mM phosphate buffer, pH 7.4) and add 100µL per well to a 96-well high-binding ELISA plate. Incubate overnight at 4℃. Wash the plate 3 times with phosphate-buffered saline containing 0.05% Tween 20 (PBST) for 5 minutes each time, then remove excess liquid by blotting with absorbent paper. Block the plate with PBST containing 1% bovine serum albumin (BSA), add 200µL per well, and incubate at room temperature for 1 hour to block nonspecific binding sites. Wash the plate 3 times with PBST for 5 minutes each time, then remove excess liquid by blotting with absorbent paper. Dilute the bacterial samples and standard solutions (known concentrations of the target protein) prepared by ultrasonic disruption in PBST containing 0.1% BSA, according to the predetermined gradient, and add 100µL per well. Incubate at room temperature for 2 hours to allow the target protein to bind to the capture antibody on the solid-phase carrier. Wash the plate 5 times with PBST for 5 minutes each time, then remove excess liquid by blotting with absorbent paper. Add the detection antibody (1:500) diluted in PBST containing 0.1% BSA to the plate, add 100µL per well, and incubate at room temperature for 1 hour to allow the detection antibody to bind to the immobilized target protein. The detection antibody is anti-human IgG antibody labeled with horseradish peroxidase (HRP), which can bind to different epitopes of the target protein. Wash the plate 5 times with PBST for 5 minutes each time, then remove excess liquid by blotting with absorbent paper. Add 100µL of TMB substrate to each well at room temperature and incubate in the dark for 15 minutes to allow HRP to catalyze the oxidation of TMB, resulting in a blue reaction product. The intensity of the reaction product is proportional to the concentration of the target protein. Add 50µL of stop solution (2M H2SO4) to each well to terminate the reaction and turn the reaction product yellow. Measure the absorbance (OD value) of each well at 450nm and plot a standard curve to calculate the concentration of the target protein in the samples [27].

### 2.20 Protein structure simulation and molecular docking

First, obtain the amino acid sequence of the target protein from the whole genome and use NCBI to perform a blast search for similar protein sequences for comparison. Then, use the multiple sequence alignment (MSA) to run the prediction script of Alphafold2 with default parameter settings, using the MSA as input, to obtain the three-dimensional structure model of the target protein, as well as the confidence score (pLDDT) and distance error (dRMSD) for each residue. Afterwards, use Swiss-model to evaluate the quality of the protein structure model predicted by Alphafold2. Through the online interface of Swiss-model, calculate the structural similarity (TM-score) and root mean square deviation (RMSD) between the predicted model and template structure, as well as the global model quality estimation (GMQE) and local model quality estimation (QMEAN) scores for the predicted model [28–29].

Subsequently, AutoDock was used to study the interaction between protein models and small molecule ligands. First, AutoDockTools was used to preprocess the predicted proteins, including removing water molecules, adding hydrogen atoms, specifying active sites, assigning atomic charges, and adjusting flexibility. Then, MGLTools was used to convert the target protein and small molecule ligands into the pdbqt format required by AutoDock, and AutoGrid was used to calculate the grid potential field of the target protein. Next, AutoDock was used for molecular docking, setting parameters such as docking iterations, population size, crossover rate, and mutation rate, and selecting the docking conformation with the highest binding affinity. PyMOL was used for visualization and analysis. Firstly, the AutoDock plugin in PyMOL was used to load the target protein and docking conformations, and the sorting and filtering functions were used to select the conformation with the lowest binding energy as the optimal conformation. Then, the display and rendering functions in PyMOL were used to adjust the colors, transparency, surfaces, and outlines of the target protein and small molecule ligands to highlight their interactions. Finally, the measurement and annotation functions in PyMOL were used to label hydrogen bonds, van der Waals forces, and hydrophobic interactions between the target protein and small molecule ligands, and the three-dimensional images of the docking results were saved [30].

### 2.21 qRT-PCR validation

After overnight culture of the strain, total RNA was extracted according to the instructions of the RNA extraction kit. The extracted bacterial RNA was used as a template for cDNA synthesis. The reverse transcription reaction was performed according to the instructions of the reverse transcription kit. 1000ng (1μg) of tRNA was prepared in a 20μl reaction system on ice. 4μl of 5× PrimeScript RT Master Mix, 1000ng of RNA, and DEPC water were added to each well of an 8-well strip. The mixture was mixed using a vortex mixer. The reaction mixture was then incubated at 37°C for 15 minutes for reverse transcription, followed by heat inactivation of the reverse transcriptase at 85°C for 5 seconds, cooling at 4°C, and storage at −20°C for later use. Using the CFX96 Real-Time system, PCR was performed in a 20μl reaction system containing 1μl of cDNA, BPGM-F/R primers at a concentration of 10μmol/L, and 10μl of 2× FastFireqPCR PreMix. The PCR program was set as follows: 95°C for 5 minutes; 40 cycles of 95°C for 5 seconds, 60°C for 30 seconds, and 72°C for 30 seconds; and a final extension at 72°C for 10 minutes. The relative expression level of the gene was calculated using the 2-ΔΔCt method with the 16SrRNA gene as the reference gene [31].

### 2.22 Coenzyme Ⅱ NADP(H) Content Assay

Cultivate two bacteria to the logarithmic growth phase. Add different concentrations of PCA to one sample for 30 minutes, and add different concentrations of 3-phosphoglyceric acid to the other sample. Add 0.5mL of acidic extraction solution and 0.5mL of alkaline extraction solution to each sample. Sonicate for 1 minute, then boil for 5 minutes, cool in an ice bath, and thaw at room temperature. Centrifuge at 10,000g for 10 minutes, transfer the supernatant to another centrifuge tube, and add equal volumes of acidic extraction solution and alkaline extraction solution. Mix well and centrifuge again for 10 minutes. Transfer the supernatant to a new centrifuge tube for measurement. Measure the absorbance at 570nm [32].

### 2.23 Reactive Oxygen Species Assay

Bacteria were cultured to the logarithmic growth phase, centrifuged at 3500rpm for 10 minutes, washed and resuspended in PBS for 3 times. Then, the fluorescent probe DCFH-DA was added and incubated on a shaker at 37℃ for 30 minutes. After that, the samples were washed and resuspended in PBS for 3 times, divided into different tubes, and different concentrations of PCA were added. The tubes were placed in a 37℃ incubator for 30 minutes. The absorbance of each sample was measured using a fluorescence microplate reader (excitation wavelength: 488nm, emission wavelength: 525nm) [33].

### 2.24 Malondialdehyde (MDA) Content Assay

Bacteria were cultured to the logarithmic growth phase, and different concentrations of PCA were added for 30 minutes. The bacteria were then washed with sterile PBS and subsequently sonicated (200w, 3s on, 10s off, repeated 30 times). In blank tubes, standard tubes, and test tubes, 0.1mL PBS, 0.1mL standard solution, and 0.1mL sample were added, respectively. Then, 0.2mL MDA detection working solution was added and mixed. The mixture was heated at 100℃ for 15 minutes, followed by cooling to room temperature in a water bath. Subsequently, 200μL was aspirated into a 96-well plate and measured at 450nm [34].

### 2.25 Intracellular ATP assay

Bacteria were cultured to the logarithmic growth phase, and different concentrations of PCA were added separately, with a negative control group set up. The cultures were placed on a shaker and incubated at 37℃ for 30 minutes. After centrifugation, the supernatant was discarded and 200μL of lysis buffer was added. The operation was carried out on ice, and then centrifuged at 4℃ and 12000g for 5 minutes. The supernatant was taken for measurement. Subsequently, 100μL of ATP detection working solution was added to a 96-well plate, followed by the addition of 20μL of sample or standard solution, with 3 replicates. Finally, the luminescence was measured using a chemiluminescence luminometer [35].

### 2.26 Statistical analysis

The results are presented as the average of three independent experiments ± standard deviation. One-way analysis of variance (ANOVA) and Student’s t-test were used to detect any significant differences between the treatments using Statplus (p<0.05) [36].

## 3 Results

### 3.1 In vitro antibacterial activity evaluation of natural plant monomer protocatechuic acid

Protocatechuic acid (PCA) is a phenolic acid compound that naturally occurs in many plants and has various biological activities. It is currently mainly used in the fields of pharmaceutical synthesis, organic intermediate synthesis, dye synthesis, and chemical reagents. It is also a core raw material for various pharmaceutical products such as Erlotinib (anti-tumor drug), Liriodenine (sodium channel inactivation gate inhibitor), Meptifan hydrochloride (respiratory system drug), Corylifolin II (treatment of hepatitis B drug), Itopride hydrochloride (novel prokinetic drug), and Yimixing (anesthetic), etc. [37]. Although PCA also has antibacterial effects, the specific mechanism of action and targets are still unclear, so it has not been widely promoted as an antibacterial product. We determined the minimum inhibitory concentration (MIC) and minimum bactericidal concentration (MBC) of two strains of carbapenem-resistant hypervirulent Klebsiella pneumoniae (CR-hvKP) isolated from animal environments. The MIC of Meropenem against MEC19116-1 and FMEC20113-2 was 64ug/mL, while the MIC and MBC of PCA against MEC19116-1 and FMEC20113-2 were both 33μM, indicating that PCA has a certain antibacterial effect against CR-hvKP. Subsequently, we used a kinetic method to determine the inhibitory effect of PCA on CR-hvKP at different time points (0, 2, 4, 6, 8, 10, 12h). The results showed (Fig.1A) that the inhibitory effect of PCA on both strains increased with increasing concentration, showing concentration dependence. At a concentration of 1×MIC, PCA completely inhibited the growth of the strains. At a concentration of 1/2MIC, PCA slowed down the growth of the bacteria.To further observe the bacterial survival count of CR-hvKP under different drug concentrations, we used DAPI staining technique for fluorescence microscopy analysis. As shown in Fig.1B, we can see that the nuclei of bacteria in the normal culture (control group) exhibit uniform blue fluorescence, indicating stable DNA content and distribution. The nuclei of bacteria treated with PCA, on the other hand, show irregular blue fluorescence, indicating damage to DNA structure and integrity. Moreover, the blue fluorescence signal significantly decreases with increasing PCA concentration. These results suggest that PCA has a bactericidal effect on CR-hvKP. To ensure the safety of PCA on the organism, we conducted a Hemolysis assay (Fig.1C) and found that the HC50 (μg/mL) of PCA on mammalian red blood cells is >20480 μg/mL, indicating that PCA is safe to use at antibacterial concentrations. In addition, we evaluated the potential development of resistance in CR-hvKP after long-term exposure to PCA. As shown in Fig.1D, after 20 generations of continuous passage at 1/2MIC concentration of PCA, the MIC remains stable, while the positive control drug tigecycline shows a 64-fold and 128-fold increase, respectively. This indicates that long-term use of the natural plant monomer PCA does not lead to a decrease in drug sensitivity of CR-hvKP, suggesting that PCA is more stable than antibiotics. Meanwhile, to evaluate whether PCA has a synergistic effect with antibiotics, we performed in vitro experiments on CR-hvKP using the checkerboard method and calculated the Fractional Inhibitory Concentration (FIC). Our experimental results (Fig.1E) show that when PCA is used in combination with meropenem, the MIC of both CR-hvKP strains is reduced by more than 4-fold. The median FIC is between 0.1 and 0.4, indicating a significant synergistic effect between PCA and meropenem. Therefore, the use of low concentrations of PCA can serve as an antibacterial adjuvant, providing a new possibility for the clinical treatment of CR-hvKP infections.

**Fig.1.**
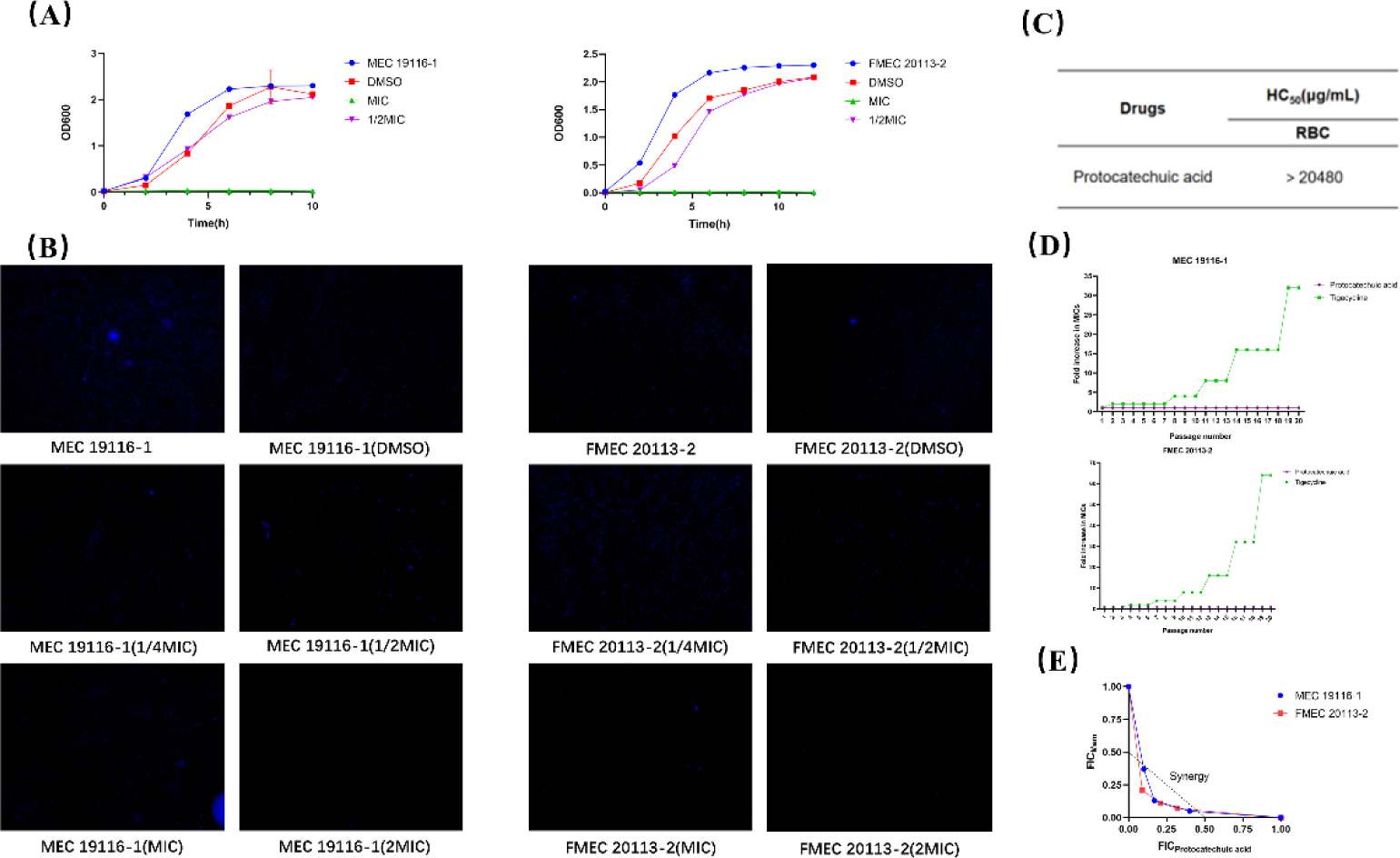
In vitro antibacterial activity evaluation of PCA. (A) Growth inhibition curve of CR-hvKP treated with PCA. (B) Bacterial DAPI staining after treatment with different concentrations of PCA. (C) HC50 (μg/mL) of PCA on mammalian red blood cells. (D) Potential development curve of drug resistance in CR-hvKP under continuous 20-generation drug treatment. (E) Synergistic study of PCA and meropenem using a checkerboard grid.

### 3.2 Inhibition of CR-hvKP biofilm by PCA at subinhibitory concentrations

Bacterial biofilms act as barriers that hinder the penetration and diffusion of antibiotics, reducing the effective concentration of antibiotics. The ability to form biofilms is closely related to the microbial physiological metabolism. In this study, we evaluated the biofilm formation of CR-hvKP treated with different concentrations of PCA. The Congo red agar plate method is a detection method that uses the principle of Congo red dye binding to the important components of biofilms, polysaccharides, to form black colonies for biofilm-positive strains and red colonies for biofilm-negative strains [38]. Using this method (Fig.2A), we observed a significant decrease in the formation of positive colonies at 1/2MIC concentration of PCA, and the extracellular secretions were also significantly lower than the control group. Subsequently, we performed crystal violet staining of CR-hvKP biofilms (Fig.2B). PCA had a weak inhibitory effect on biofilms at low concentrations, but as the drug dosage increased, PCA showed a significant inhibitory effect on CR-hvKP biofilms (p<0.01), especially with an average inhibition rate of over 60% for MEC19116-1. EPS (extracellular polymeric substances) are high molecular weight natural polymers secreted by microorganisms into their environment, which have important effects on bacterial virulence, biofilm formation, and stability. The main components are polysaccharides, proteins, DNA, lipids, and other macromolecules [39]. Through the extraction and measurement of esp after PCA treatment (Fig.2C), it was found that PCA did not exhibit dose-dependent inhibition of eps at low concentrations, but instead caused compensatory increase. However, when the drug concentration exceeded a certain threshold, it could significantly inhibit esp (p<0.001). Overall, the inhibition of eps was consistent with the results of crystal violet staining. In addition, during the process of extracting esp from bacteria using a 6-well plate, we also found that the bacteria in the 6-well plate showed different color changes at different concentrations of PCA (Fig.2D) at 1/8MIC, 1/4MIC, 1/2MIC, MIC, and 2MIC drug concentrations. This may be due to the impact of PCA on bacterial metabolism [40].

**Fig.2.**
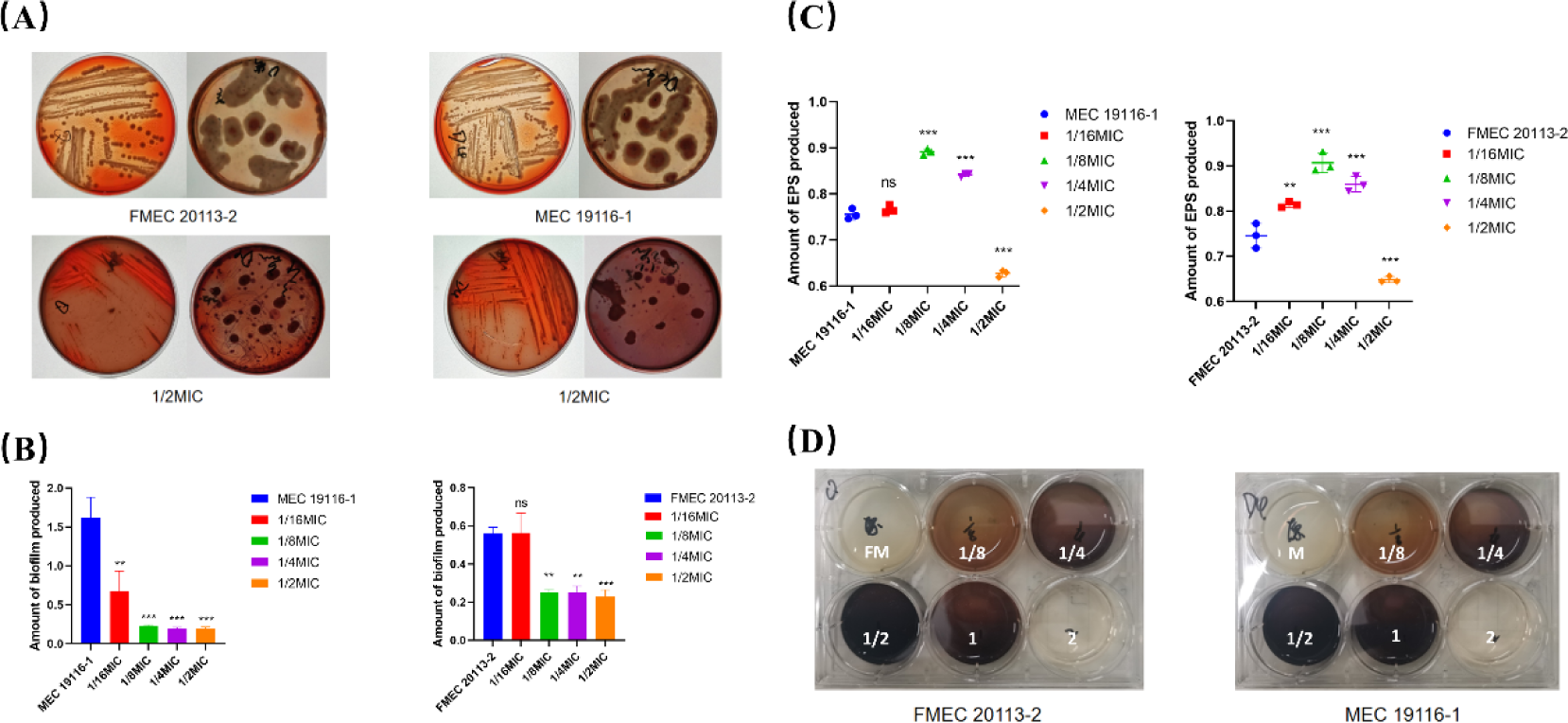
Evaluation of the inhibitory ability of sub-inhibitory concentration of PCA on CR-hvKP biofilm. (A) Observation of bacterial biofilm in control group and sub-inhibitory concentration PCA treatment group. (B) Crystal violet staining of biofilm after treatment with different concentrations of PCA under sub-inhibitory conditions. (C) Measurement of extracellular polymeric substances. (EPS) after treatment with different concentrations of PCA under sub-inhibitory conditions. (D) Color change of bacterial culture medium under different concentrations of PCA treatment (numbers in the figure represent drug concentrations). Values are presented as mean ± SD. ns represents (P>0.05), * represents (P<0.05), ** represents (P<0.01), *** represents (P<0.001), all compared to the control group.

### 3.3 The effect of subinhibitory concentration of PCA on the cell membrane of CR-hvKP

The bacterial cell membrane is the first line of defense against antibiotic resistance in bacteria. Bacteria can reduce the entry of antibiotics by changing the permeability of the cell membrane, thereby developing resistance. This change can be a defect or reduction in the expression of pore proteins, or a change in the membrane potential [41]. In addition, the relationship between the bacterial cell membrane and metabolism is a complex and interesting topic. Due to the rich enzyme system present on the membrane, it serves as a site for many important metabolic activities in bacteria [42]. The cell membrane is also closely related to bacterial quorum sensing and biofilm formation. Quorum sensing signal molecules freely diffuse or actively transport through the bacterial cell membrane, bind to receptors inside and outside the cell, and trigger downstream signaling pathways as well as the formation and maintenance of biofilms [43–44]. In this study, we first observed the morphology of the bacterial cell membrane after treatment with sub-inhibitory concentration of PCA using scanning electron microscopy (Fig.3A). We found that the cell membrane of FMEC20113-2 became rough and uneven under PCA treatment, while the control group maintained a smooth and uniform morphology. The cell membrane of MEC19116-1 even showed damage and leakage of cellular contents. This indicates that PCA has a significant damaging effect on the cell membrane of CR-hvKP and affects membrane permeability. The permeability of the bacterial cell membrane has a bidirectional impact on bacterial metabolism. Normal metabolic activities require stable membrane permeability to ensure the exchange of substances between the inside and outside of the cell and the stability of the internal environment. On the other hand, bacterial metabolism can also affect changes in cell membrane permeability by altering the composition and structure of the cell membrane or producing substances that affect membrane permeability [45]. Therefore, in the following study, we evaluated the inner membrane permeability using the non-permeable nucleic acid binding dye PI, which can only enter the cell through a damaged cell membrane and bind to DNA or RNA, emitting red fluorescence [46]. By measuring the fluorescence intensity (Fig.3B), we found that the fluorescence value after PCA treatment increased with the drug concentration, indicating a dose-dependent positive correlation between inner membrane permeability and PCA (P<0.01). Furthermore, we used a fluorescent dye NPN (1-N-phenylnaphthylamine) to measure the outer membrane permeability of CR-hvKP. NPN can dissolve in the lipid bilayer but cannot penetrate intact bacterial outer membranes. When the bacterial outer membrane is damaged, NPN can enter the periplasmic space of the bacteria and interact with hydrophobic molecules, emitting fluorescence, reflecting changes in bacterial outer membrane permeability [47]. As shown in (Fig.3C), the fluorescence intensity after PCA treatment increased with the drug concentration, indicating a dose-dependent positive correlation between outer membrane permeability and PCA (P<0.001). As for why the fluorescence intensity decreased at MIC concentration, we believe that PCA at MIC concentration caused obvious damage to the outer membrane of the bacteria, causing it to detach from the cell wall, and the hydrophobic environment of the periplasmic space no longer existed, resulting in a decrease in fluorescence intensity [48]. Changes in membrane permeability also affect membrane potential. Membrane potential is a crucial source of free energy for bacteria and can affect bacterial signal transduction, stress regulation, and antibiotic resistance. For example, the bacterial respiratory chain and ATP synthase are located on the cell membrane, and they require membrane permeability to maintain membrane potential and proton gradient for energy conversion and utilization [41]. In this study, we measured the Δψ of the cell membrane using the fluorescence DiSC3(5) method [49]. DiSC3(5) is a membrane potential-sensitive probe that aggregates in the phospholipid bilayer and causes self-quenching of the dye. When the membrane is depolarized, the potential dissipates, and diSC3(5) is released into the solution, causing an increase in fluorescence, which is proportional to the degree of membrane potential decrease [50]. As shown in (Fig.3D), the fluorescence intensity increased with the drug concentration after PCA treatment (P<0.05), indicating that the Δψ depolarization of the CR-hvKP cell membrane is affected by PCA. In summary, the results of the cell membrane study further illustrate the impact on bacterial activity and metabolism.

**Fig.3.**
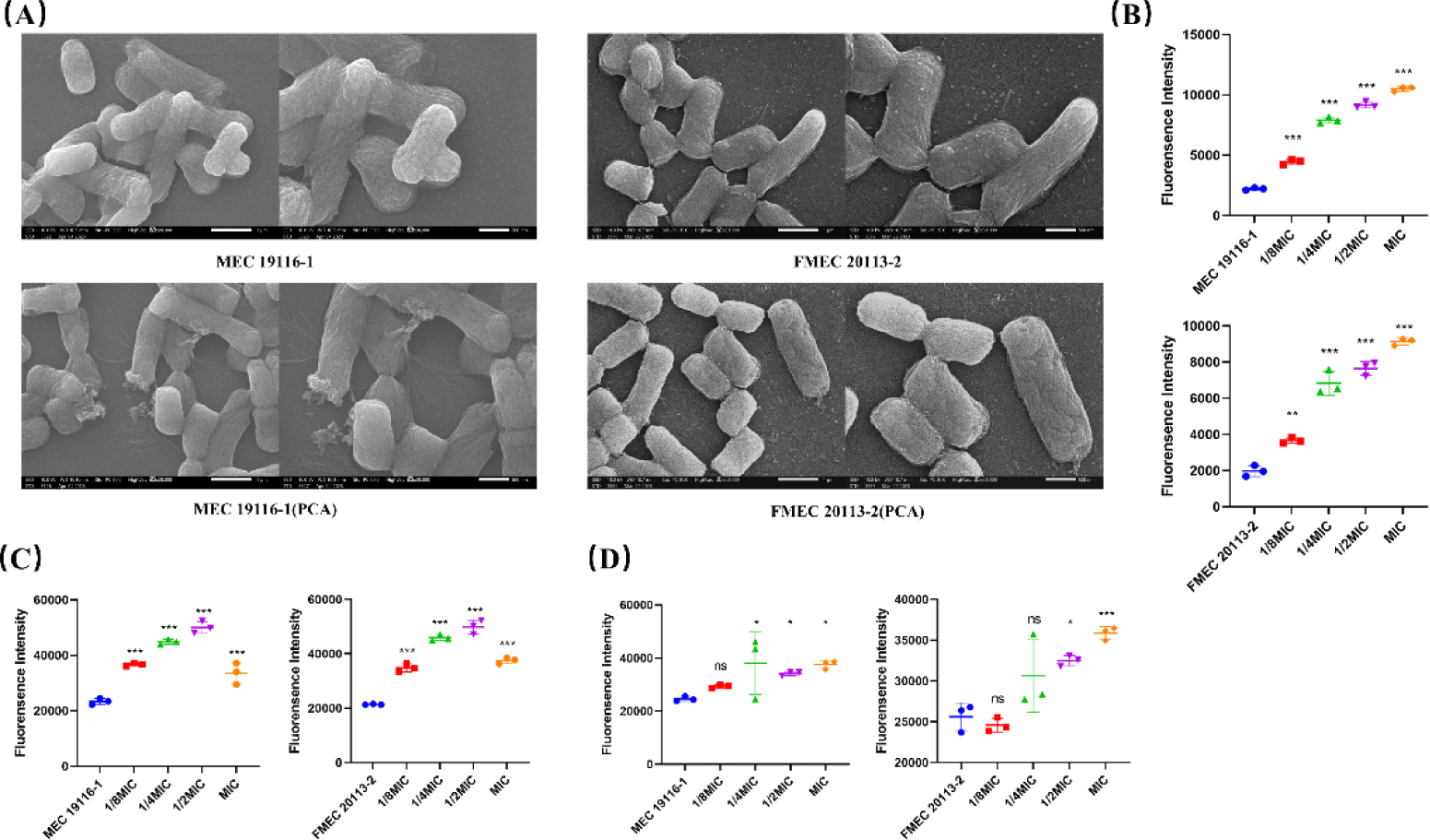
The effect of subinhibitory concentrations of PCA on the bacterial cell membrane of CR-hvKP as studied by PCA. (A) Morphological observation of bacteria in the control group and PCA treatment group. (B) Intracellular membrane permeability after treatment with different concentrations of PCA under subinhibitory conditions. (C) Extracellular membrane permeability after treatment with different concentrations of PCA under subinhibitory conditions. (D) Cell membrane Δψ after treatment with different concentrations of PCA under subinhibitory conditions. Values are presented as mean ± SD. In the figure, ns indicates (P>0.05), * indicates (P<0.05), ** indicates (P<0.01), *** indicates (P<0.001), all compared to the control group.

### 3.4 The effect of subinhibitory concentration of PCA on bacterial metabolism

As mentioned above, the steady state of the cell membrane has a bidirectional effect on bacterial metabolism. In order to study the effect of PCA on bacterial metabolic status, we used AlamarBlue to detect the bacterial viability and metabolic activity after drug treatment. During normal metabolic processes, the intracellular environment of bacteria changes from an oxidative environment to a reducing environment. The active component of AlamarBlue, resazurin, is a non-toxic and membrane-permeable blue dye. After being taken up by the cells, it is reduced by metabolic intermediates, and the reduced product, resorufin, appears pink and exhibits strong fluorescence [51]. In this study (Fig.4A), we found that the fluorescence intensity of bacteria decreased after PCA treatment (P<0.05). With increasing drug concentration, a color different from pink was observed, indicating that the drug reduced the metabolic activity and reducing ability of the bacteria.

**Fig.4.**
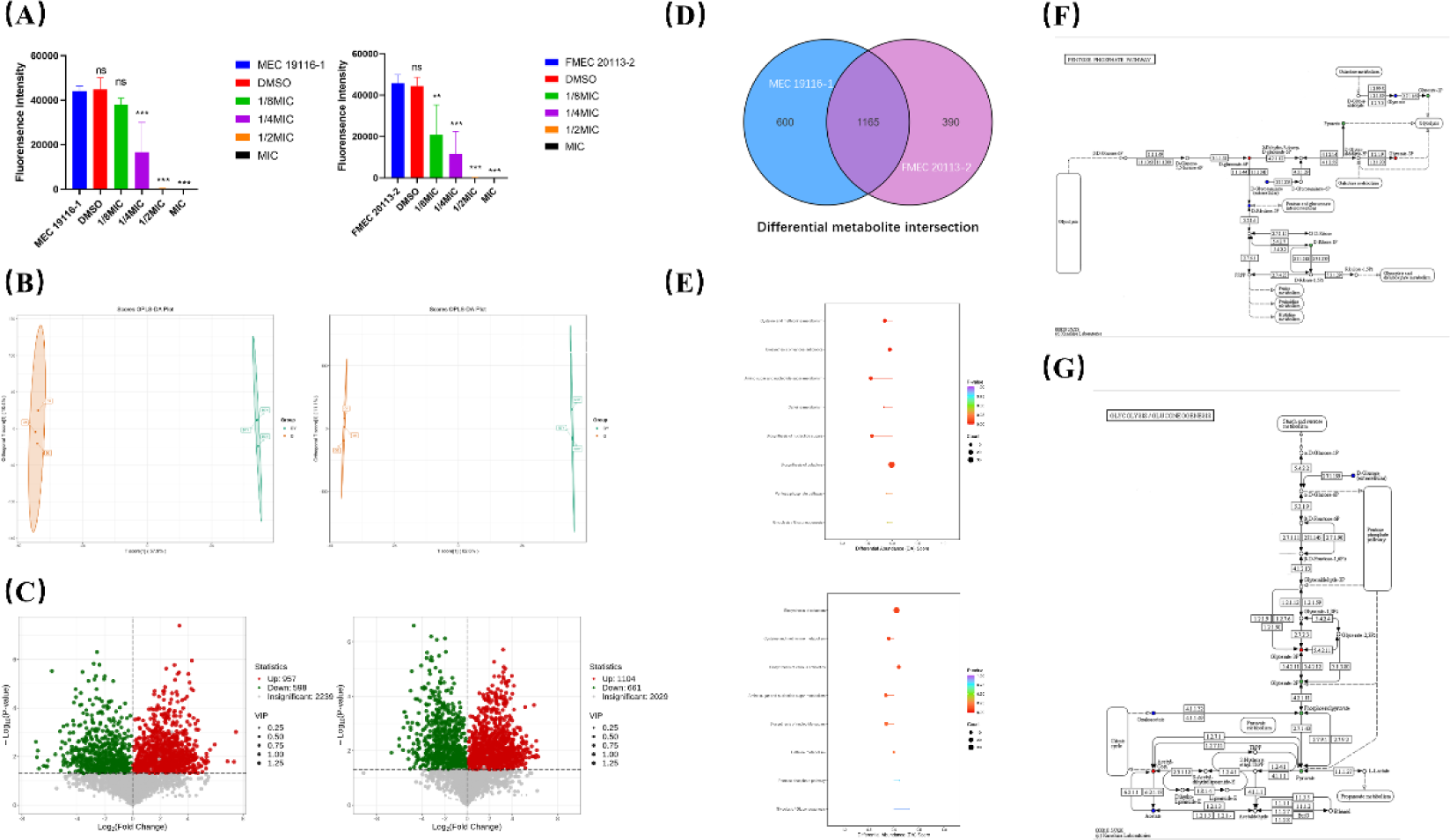
The effect of subinhibitory concentrations of PCA on bacterial metabolism as determined by PCA. (A) AlamarBlue assay measuring the metabolic activity of CR-hvKP after PCA treatment. (B) Differential metabolite profiles after PCA treatment at subinhibitory concentrations (P<0.05). (C) Volcano plot of differential metabolites; significantly upregulated and downregulated metabolites are shown in red and green, respectively, while non-significant differentially expressed metabolites are shown in gray. (D) Venn diagram of differential metabolites between the two groups. (E) KEGG enrichment analysis of differential metabolites, showing the most significantly enriched pathways. (F) Changes in the pentose phosphate pathway and associated metabolites; significantly upregulated and downregulated metabolites are shown in red and green, respectively. (G) Changes in the glycolysis pathway and associated metabolites; significantly upregulated and downregulated metabolites are shown in red and green, respectively.

Subsequently, we used metabolomics to analyze the bacterial metabolic activity of CR-hvKP treated with PCA, in order to gain a more comprehensive understanding of how PCA affects CR-hvKP and its underlying mechanisms. Metabolomic analysis was performed using a UPLC-Q-TOF/MS system and multivariate statistical analysis to analyze the perturbation of metabolite profiles in the cells of two CR-hvKP strains treated with PCA. Differential metabolites (DMs) were selected based on the OPLS-DA model, which provided convincing parameters with high R2Y and Q2Y values. Both the control group and the treatment group showed distinct clustering behavior. All replicates of each group were assigned to the same cluster, confirming the observed differences in metabolite profiles between the control and treatment groups (Fig.4B).As shown in Fig. 4C, in the FMEC20113-2 group, compared to the control group, 2029 significantly altered metabolites were identified, including 1104 upregulated and 661 downregulated. Similarly, in the MEC19116-1 group, compared to the control group, 2239 significantly altered metabolites (VIP>1, P<0.05) were identified, including 957 upregulated and 598 downregulated. Meanwhile, we presented the shared differential metabolites of the two bacterial groups in a Venn diagram (Fig. 4D).

Finally, we performed KEGG enrichment analysis to show the significantly enriched pathways in two groups of bacteria after PCA treatment, and analyzed the overall changes in these metabolic pathways based on differential abundance scores (Fig. 4E and F). We found that most of these co-enriched metabolic pathways showed downregulation. Through analysis, we found that the metabolic pathways including “Cysteine and methionine metabolism”, “Biosynthesis of various antibiotics”, “Aminosugar and nucleotide sugar metabolism”, “Caffeine metabolism”, “Biosynthesis of nucleotide sugars”, and “Biosynthesis of cofactors” were strongly associated with the Pentose phosphate pathway (PPP). For example, in the pathways of “Biosynthesis of nucleotide sugars” and “Aminosugar and nucleotide sugar metabolism”, bacteria can generate ribose-5-phosphate from glucose or other hexose sugars through the PPP pathway, and then convert it into various aminosugars and nucleotide sugars through different transferases [52–54]. In the pathway of “Cysteine and methionine metabolism”, bacteria reduce inorganic sulfur to hydrogen sulfide using NADPH provided by the PPP pathway, and then react with serine or methionine to generate cysteine or methionine [55]. In addition, other significant metabolic pathways were related to amino acid metabolism, which requires amino acids as precursors and depends on ribose-5-phosphate produced by the PPP pathway [54]. In summary, the PPP pathway is a key influencing pathway for drug response, as it is closely related to other significantly changed bacterial metabolic pathways. It can provide important metabolic intermediates such as ribose-5-phosphate and NADPH, and participate in the synthesis and regulation of various biomolecules in these pathways. When the PPP pathway is affected, these associated metabolic pathways show downregulation.

Therefore, we analyzed the changes in the metabolites of the pentose phosphate pathway (PPP) in two groups of bacteria after PCA treatment. The PPP, also known as the pentose phosphate pathway, is a metabolic pathway in bacteria that can produce ribose-5-phosphate and NADPH in cells, which are essential for the synthesis of biomolecules such as nucleic acids and coenzymes [54]. Moreover, the PPP affects the energy metabolism of bacteria by regulating glycolysis and the citric acid cycle [56]. The PPP consists of two stages: oxidative and non-oxidative. In (Fig.4G), it can be seen that the oxidative stage of the PPP starts with β-D-glucose-6-phosphate, which is an intermediate product of glycolysis and the first substrate of the PPP. It is oxidized by glucose-6-phosphate dehydrogenase (G6PDH) to D-glucono-1,5-lactone-6P, while reducing NADP+ to NADPH. This is the first reaction of the PPP and the rate-limiting reaction, which determines the activity of the PPP. D-glucono-1,5-lactone-6P is hydrolyzed by a hydrolase to D-gluconate-6P, also known as 6-phosphogluconate, which is an intermediate product of the PPP. 6-phosphogluconate is oxidized by 6-phosphogluconate dehydrogenase (6PGDH) to D-ribulose-5P, while reducing NADP+ to NADPH. This is the second reaction of the oxidative stage of the PPP and the last reaction of the oxidative stage, which produces two NADPH and one 5-phosphoribose. The non-oxidative stage of the PPP starts with D-ribulose-5P in the top right corner, which is the last product of the oxidative stage and the first substrate of the non-oxidative stage [56]. It is isomerized to D-ribose-5P by ribose-5-phosphate isomerase (RPI). D-ribose-5P is an important carbon skeleton that can participate in various biological processes such as the synthesis of nucleic acids and coenzymes, as well as carbon metabolism [57].

Through metabolomics analysis (Fig.4G), it was found that the glycerate-3P in bacteria showed a significant upregulation after treatment with both drugs. Glycerate-3P is a common molecule in biological cells and is an intermediate product of the glycolysis (EMP) pathway. It is usually used as a raw material for the synthesis of other organic molecules such as glucose, nucleic acids, and fatty acids [58]. When the concentration of glycerate-3P is too high, it can have a certain impact on the bacteria’s PPP pathway. This is because excessive glycerate-3P can reduce the flux of the PPP pathway. The excessive accumulation of this intermediate product will compensatorily change the activity and expression of the rate-limiting enzyme in the pathway, which is glucose-6-phosphate dehydrogenase (G6PDH) in the PPP pathway. This compensatory mechanism eventually evolves into a stress response, where G6P is blocked from entering the reaction for a long time, severely slowing down the oxidative stage of the PPP pathway, resulting in a decrease in NADPH production, an increase in intracellular oxidative stress, and damage to DNA, proteins, and lipids, ultimately leading to apoptosis or necrosis [59]. NADPH is an important reducing agent that participates in biosynthesis and antioxidant processes. On the other hand, due to the slowdown of the oxidative stage of the PPP, the generation of ribose-5-phosphate will decrease, further affecting the synthesis of nucleic acids and coenzymes, inhibiting bacterial growth and metabolism [57]. For example, as observed in (Fig.4G), the significant downregulation of D-ribose-1P, which is converted from D-ribose-5P (a type of 5-phosphoribose) catalyzed by ribose-phosphate isomerase (RPE), is a result of the inhibition of the PPP pathway.

We speculate that pyruvic acid (PCA), as an organic acid, can enter bacterial cells and affect the bacterial EMP, thereby inhibiting bacterial growth and metabolism. In glycolysis (Fig.4H), glucose is converted to glyceraldehyde-3-phosphate (G3P), which is then phosphorylated and converted to 1,3-diphosphoglycerate (1,3-DPG). Subsequently, under the action of phosphoglycerate kinase (PGK), 1,3-DPG is converted to glycerate-3-phosphate (G3P), and then under the action of phosphoglycerate mutase (PGAM), G3P is converted to glycerate-2-phosphate (G2P), finally leading to the production of pyruvate [58]. It is worth noting that the reactions in this stage are reversible and adjusted by the relative concentration of intermediate products, so G3P can be converted between different pathways of EMP to maintain normal metabolism [58]. Therefore, if PCA inhibits the activity of PGAM, G3P cannot be converted to G2P, and even G2P may be downregulated. However, in this case, bisphosphoglycerate mutase (BPGM) can catalyze the conversion of G3P to glycerate-2,3-phosphate (G2,3P), thereby continuing to participate in the metabolic process and ensuring the steady progress of EMP. This is a compensatory mechanism of bacteria. Although the gene expression level of BPGM in Klebsiella pneumoniae is relatively low and only induced when the pathway is abnormal, it can alleviate the toxic effects caused by the excessive accumulation of G3P [60–61]. However, when BPGM is also inhibited by PCA, the concentration of G3P will continue to rise because it cannot be effectively utilized or cleared. This further leads to the blockage of the EMP pathway itself and the pentose phosphate pathway, resulting in metabolic disorders.

The mainstream metabolic pathway in most microorganisms is the EMP pathway, but few microorganisms rely solely on the EMP pathway for metabolism, mostly through the combined action of the EMP and PPP pathways. Based on the analysis of metabolomics reports, we found that PCA mainly inhibits the action of EMP and simultaneously blocks the PPP pathway in CR-hvKP, leading to an imbalance in the redox state of the bacterial body and disruption of energy metabolism, as well as inhibition of biofilm function and changes in membrane permeability.

### 3.5 Changes in key metabolic enzymes

To validate the results obtained in metabolism, we first detected the activity of G6PDH after PCA treatment of bacterial cells, as shown in (Fig.5A). PCA stimulated bacterial compensation at low concentrations, leading to an increase in G6PDH activity. In this case, the flux of the pentose phosphate pathway would increase, the NADPH/NADP+ ratio would rise, and thus meet the requirements for cellular biosynthesis and resistance to oxidative stress under drug stimulation. However, as the concentration of PCA increased, this compensatory mechanism could not be maintained, resulting in a decrease in G6PDH activity (P<0.05). At the same time, we added exogenous glycerate-3P and retested the activity of G6PDH, as shown in (Fig.5B). When 0.2μM and 0.5μM were added, the activity of G6PDH decreased to half of the original level. However, when we added >1μM glycerate-3P, G6PDH activity decreased significantly (P<0.001). This indicates that glycerate-3P does indeed inhibit the function of G6PDH under conditions of excessive accumulation, as we expected.

**Fig.5.**
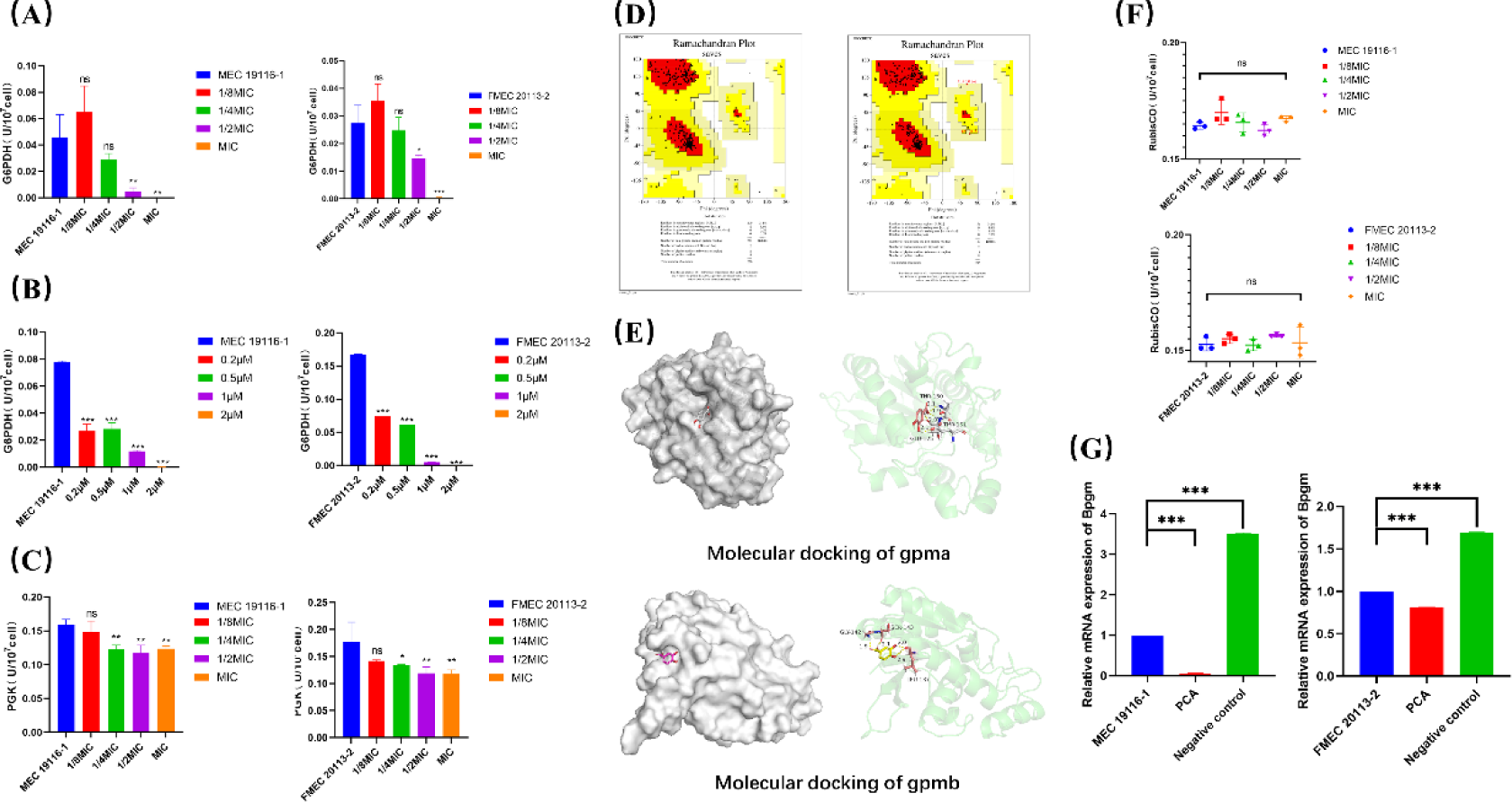
The effect of subinhibitory concentration of PCA on key metabolic enzymes. (A) Effect of subinhibitory concentration of PCA on G6PDH activity. (B) Effect of exogenously added glycerate-3P at different concentrations on G6PDH activity. (C) Effect of PCA on PGAM enzyme activity. (D) Quality assessment of 3D structure modeling of gpma and gpmb proteins. (E) Molecular docking simulation of PCA with gpma and. (F) Effect of PCA on BPGM enzyme activity. (G) Transcriptional effect of PCA on BPGM at subinhibitory concentration. Values are presented as mean ± SD. In the figure, ns indicates (P>0.05), * indicates (P<0.05), ** indicates (P<0.01), *** indicates (P<0.001), all compared to the control group.

Meanwhile, we also detected the activity of PGAM enzyme through ELISA, and the results are shown in (Fig.5C). As the concentration of PCA increases, the activity of PGAM decreases, indicating that PCA indeed has an inhibitory effect on PGAM activity, and this effect shows a dose-dependent relationship (P < 0.05). In order to explore the interaction between PCA and PGAM, we performed molecular docking between PCA and PGAM. In Klebsiella pneumoniae, PGAM is mainly encoded by gpma and gpmb. Although the proteins encoded by gpma and gpmb have the same enzymatic activity, they have different functional domains, so we docked both of them. In our previous study, we obtained the whole genome sequence of FMEC20113-2 and obtained the amino acid sequence information of gpma and gpmb from it. Then we simulated their protein structures using Alphafold2 and evaluated the quality of protein structure modeling using Swiss-model (Fig.5D) to obtain the most suitable protein structure. Based on this, we employ Autodock molecular docking software to integrate PCA with proteins, discernring that both PCA and the proteins encoded by gpma and gpmb exhibit robust affinity for attachment.. Finally, we used pymol software to display the key amino acids of the two proteins binding with PCA and analyzed their interactions. As shown in (Fig.5E), PCA exerts its inhibitory effect on activity by binding to the amino acid residues GLU-125, THR150, and THR151 of gpma, and by binding to the amino acid residues LEU337, GLY342, and SER343 of gpmb. The main interaction between them is through hydrogen bonding, which affects the function of the target protein.

In the previous discussion, we mentioned that bacteria can convert excessive glucerate-3P through compensatory expression of BPGM. Therefore, in this study, we simultaneously detected the activity and content of BPGM enzyme, as shown in (Fig.5F). Although the activity of BPGM fluctuated slightly under the influence of the drug, it remained at a low level of activity regardless of the drug concentration (P>0.05). Therefore, PCA may not have a significant impact on the activity of BPGM. Since the expression level of the BPGM gene itself is low in microorganisms, we suspect that the inhibitory effect of PCA occurs during its transcription process. Therefore, we conducted RT-qPCR experiments (Fig.5G) and found that compared to the negative group treated with DMSO, PCA significantly downregulated the transcription of bpgm, and the inhibitory effect of PCA was more pronounced in the MEC19116-1 group (P<0.001). As for the upregulation observed in the DMSO group, this may be due to bacterial compensation caused by environmental changes.

In conclusion, as we obtained in the metabolomics analysis, the upregulation of the signature metabolite glycerate-3P is due to the inhibition of the EMP pathway. The accumulation of this intermediate product will compensatorily affect the rate-limiting enzyme of the PPP pathway, ultimately leading to a decrease in the activity and expression of G6PDH, resulting in a significant inhibition of the PPP metabolic pathway and causing endogenous oxidative stress.

### 3.6 Endogenous Oxidative Stress in Bacteria

The pentose phosphate pathway (PPP) is a key pathway for regulating the NADPH/NADP+ balance, ensuring the redox balance inside bacterial cells. In addition, NADPH, as an important reducing agent, can provide reducing power to drive various cellular synthesis reactions. For example, NADPH can participate in fatty acid synthesis by extending the fatty acyl chain through the reduction of fatty acyl-ACP reductase (FASR). NADPH can also participate in cholesterol synthesis by generating methylglutaconyl-CoA through the reduction of HMG-CoA reductase (HMGR). NADPH is also involved in nucleotide synthesis by generating deoxyribonucleotides through the reduction of nucleotide reductase (NR). NADPH can also participate in the synthesis of amino acids such as proline, tryptophan, and tyrosine. NADPH is also an important signaling molecule that can affect cell signaling by regulating intracellular calcium levels, nitric oxide synthase (NOS) activity, and oxidative stress response. Some studies have shown that NADPH can regulate biofilm formation by affecting the c-di-GMP levels inside bacteria. c-di-GMP is a widely distributed second messenger in bacteria that can regulate bacterial motility, adhesion, and biofilm formation. Additionally, NADPH is also an important antioxidant that can eliminate reactive oxygen species (ROS) by reducing molecules such as glutathione (GSH) and thioredoxin (Trx). Therefore, when the PPP pathway is inhibited, NADPH is downregulated, resulting in a decrease in the bacteria’s antioxidant capacity and susceptibility to oxidative damage. In this study, we measured the intracellular NADPH/NADP+ ratio after PCA treatment, as shown in Figure 6A. When PCA was used at a concentration below 1/2MIC, the NADPH/NADP+ ratio significantly decreased. This is because the inhibition of glucose-6-phosphate dehydrogenase (G6PDH) prevents the reduction of NADP+ to NADPH. As the drug concentration increases, the synthesis of NADP+ is also affected, for example, by inhibiting the activity of NAD kinases (NADKs), leading to a decrease in NAD+ conversion to NADP+. In this case, the NAD+ content also decreases, resulting in an increase in the NADPH/NADP+ ratio. Therefore, combined with previous studies such as the significant downregulation of amino acid and nucleotide synthesis in metabolomics, as well as the inhibition of biofilm formation, it is evident that the alteration of the NADPH/NADP+ balance severely affects the growth and function of CR-hvKP.

Based on this, we also determined the levels of bacterial ROS and malondialdehyde (MDA). ROS are a type of highly reactive free radicals that can damage bacterial structures such as membranes, proteins, and nucleic acids, leading to bacterial dysfunction and death [66]. MDA is the main product of lipid peroxidation and a commonly used indicator to measure the degree of lipid oxidative damage. When the intracellular ROS levels are high, unsaturated fatty acids react with ROS to produce a series of peroxides and MDA as end products [67]. The results, as shown in (Fig. 6B and C), indicate a positive correlation between the concentration of PCA and the levels of bacterial intracellular ROS and MDA (P < 0.001). This result is consistent with the changes in NADPH/NADP+ and further supports the notion that the disruption of bacterial PPP metabolism is the cause of bacterial oxidative stress.

Both oxidative stress-induced damage to the respiratory chain and changes in cell membrane potential, as well as inhibition of the EMP pathway, can reduce the decline of intracellular ATP, affecting energy balance and bacterial function. Therefore, we also measured the changes in intracellular ATP levels, as shown in (Fig.6D). At low concentrations, PCA does not affect the intracellular ATP levels, but instead increases its content. This is due to the compensatory effect of low-dose drug stimulation on bacteria. However, as the concentration increases, due to the destructive and inhibitory effects of PCA on the respiratory chain and EMP, intracellular ATP levels decrease.

**Fig6.**
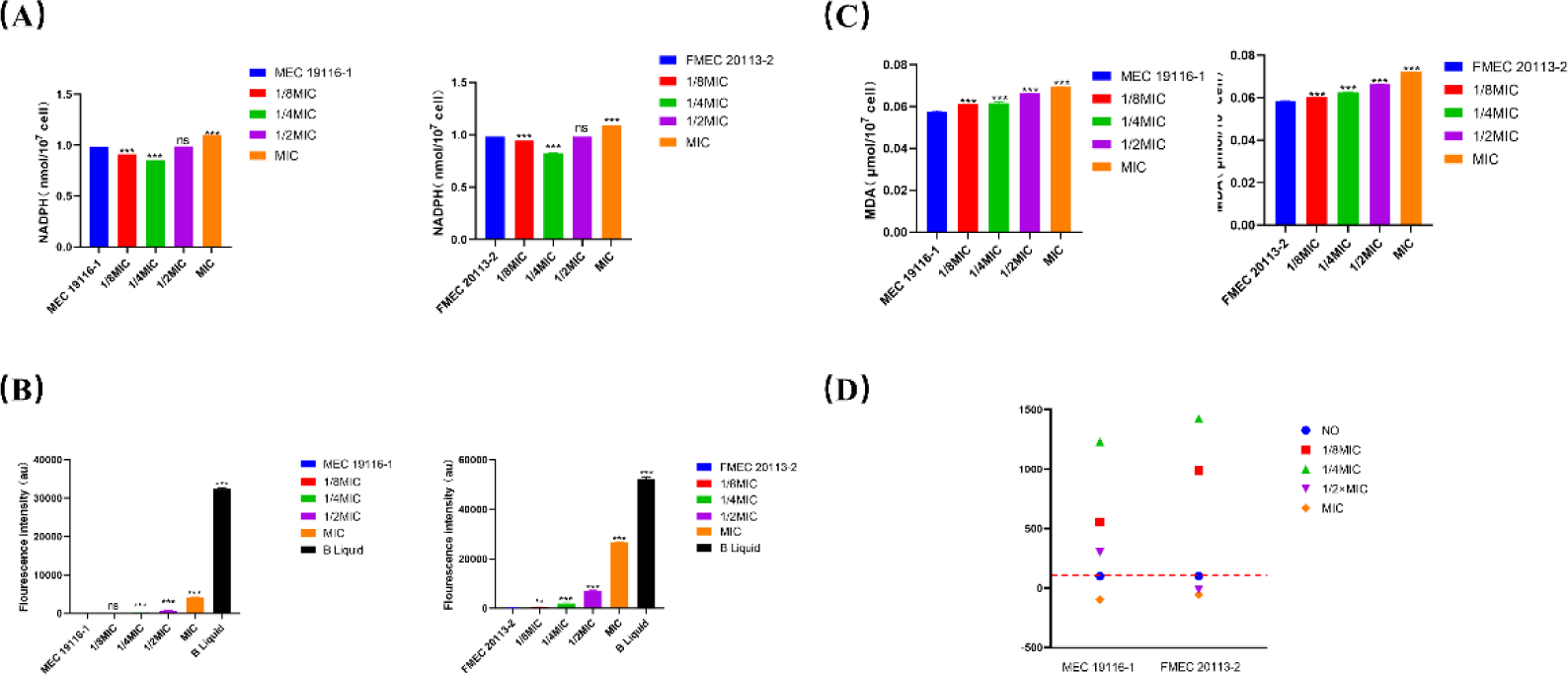
The effect of subinhibitory concentration of PCA on the redox state of bacteria as analyzed by PCA. (A) The effect of subinhibitory concentration of PCA on intracellular NADPH/NADP+ ratio. (B) The effect of subinhibitory concentration of PCA on intracellular ROS levels. (C) The effect of subinhibitory concentration of PCA on intracellular MDA levels. (D) The effect of subinhibitory concentration of PCA on intracellular ATP levels.Values are presented as mean ± SD. In the figure, ns indicates (P>0.05), * indicates (P<0.05), ** indicates (P<0.01), *** indicates (P<0.001), all values compared to the control group.

## 4. Discussion

In the context of the widespread spread of drug-resistant bacteria, the treatment of antibiotics has become difficult, and increasing the dosage can lead to a vicious cycle, with bacteria becoming less and less sensitive to antibiotics or even developing new resistance [68]. People have begun to explore the possibility of using bioactive molecules from natural plant sources as antibacterial drugs and antibacterial enhancers [69–70]. Protocatechuic acid is a polyphenolic compound found in various natural plants. Previous studies have focused on its excellent antioxidant properties, as well as its anti-inflammatory and anticancer effects in the body [71–72]. Considering that PCA is an important raw material that has been used in pharmaceutical synthesis for a long time and has a high level of safety, this study investigated the antibacterial activity of PCA against drug-resistant bacteria and its related mechanisms. Based on antibacterial tests, we found that PCA can inhibit the growth of CR-hvKP at lower concentrations. In addition, PCA also has a synergistic effect with the carbapenem antibiotic meropenem. Continuous treatment with PCA does not induce resistance in CR-hvKP compared to other antibiotics. Since subinhibitory concentrations of PCA can synergistically inhibit bacteria with meropenem, we also studied the inhibitory effect of PCA on CR-hvKP biofilms and its impact on cell membranes at subinhibitory concentrations. The results showed that besides being used as an antibacterial drug on its own, PCA can also act as an antibacterial enhancer by inhibiting biofilm formation and altering membrane permeability at low concentrations, making it a potential candidate drug against drug-resistant bacteria.

The metabolic homeostasis of the internal environment is crucial for the function and growth of bacteria [73]. Key enzymes in metabolic pathways and secondary metabolites are promising targets for screening candidate antibacterial agents [74]. For example, research has targeted key enzymes in the bacterial folate metabolism pathway to design folate antagonist antibacterial drugs, such as inhibitors of dihydrofolate reductase (DHFR) [75]. There are also studies that use metabolomics-based methods to explore the interaction mechanisms between antibiotics and bacteria, thereby screening for more novel antibiotics [76]. Due to the antibacterial activity of certain secondary metabolites, such as pyocyanin (PYO) in Pseudomonas aeruginosa and pyocyanin in Staphylococcus aureus [77], we can also develop antibacterial drug adjuvants targeting the metabolites themselves. Since there is a bidirectional influence between bacterial cell membrane and bacterial metabolic activity [42][45], the function of biofilms is also regulated by bacterial metabolism [78–79]. Therefore, to gain a deeper understanding of the biological mechanism of PCA on CR-hvKP, metabolomics methods were used to describe the major changes in the biological processes of CR-hvKP after PCA treatment. The results showed that PCA interfered with multiple biological pathways related to amino acid metabolism in CR-hvKP, considering that the production of amino acids mainly relies on the precursors provided by the PPP pathway. We speculate that the main metabolic pathways disrupted by PCA are the PPP pathway and the closely related glycolysis pathway.

The main pathway for microbial energy metabolism is the glycolysis pathway, in which bacteria break down sugars into pyruvic acid or other organic acids and generate energy [80]. There are four main pathways for bacterial glycolysis: EMP, HMP, ED, and WD. Most microorganisms rely on the combined action of EMP and HMP [81]. The EMP pathway is the most common glycolysis pathway and the first stage of cellular respiration. It consists of 10 steps, converting 1 molecule of glucose into 2 molecules of pyruvic acid, and producing 2 molecules of ATP and 2 molecules of NADH [82]. The HMP pathway, also known as the PPP pathway mentioned earlier, is a pentose phosphate pathway. It degrades 6-phosphogluconate as a substrate and is divided into two stages: oxidative and non-oxidative stages. This pathway can provide reducing power NADPH and carbon skeletons for biosynthesis, and can also connect with the EMP pathway to expand the range of carbon sources utilization [83]. Studies have shown that EMP and PPP are interdependent and mutually influenced. For example, the product of glycolysis, pyruvic acid, can enter the tricarboxylic acid cycle, and the intermediate products of the tricarboxylic acid cycle can enter the pentose phosphate pathway. They collectively generate energy, reducing power, and precursor substances in the cytoplasm, and regulate their own activity and flow according to the needs of the cytoplasm and changes in the environment [84]. The metabolomics results of this study indicate that the metabolite 3-phosphoglyceric acid in the EMP pathway is significantly upregulated. The abnormal accumulation of this intermediate metabolite may lead to blockage of the PPP pathway, resulting in a series of cellular functional disorders. For example, the accumulation inhibits downstream reactions, reduces the production of NADPH, leading to oxidative-reductive imbalance in the cell; reduces the production of R5P, affecting nucleic acid synthesis; reduces the production of GAP, reducing energy supply, etc. [85–87]. In this study, we found that the activity of the rate-limiting enzyme G6PDH in the PPP pathway is affected by the intermediate metabolite 3-phosphoglyceric acid, and the metabolic disorder of 3-phosphoglyceric acid is the result of the inhibition of the key enzyme PGAM by PCA. We verified this result through molecular docking. In addition, when 3-phosphoglyceric acid is metabolically abnormal, microorganisms compensatorily express BPGM to alleviate this situation [88]. We found through enzyme-linked immunosorbent assay and real-time fluorescence quantitative PCR that PCA significantly downregulates the expression of BPGM. These results indicate that PCA inhibits protein activity by hydrogen bonding with PGAM amino acid residues, and also inhibits the expression of the bypass pathway BPGM, leading to abnormal accumulation of the key metabolite 3-phosphoglyceric acid. This accumulation not only affects the normal operation of the EMP pathway but also negatively inhibits the PPP pathway, ultimately leading to the comprehensive shutdown of microbial energy metabolism.

It is worth noting that the inhibition of the pentose phosphate pathway (PPP) alters the redox state of bacteria, leading to endogenous oxidative stress. This is because the PPP is the main source of NADPH, and the balance between NADPH and NADP+ is crucial for resisting reactive oxygen species (ROS) [89]. Inhibition of the PPP reduces the level of NADPH, making bacteria more susceptible to ROS damage and disrupting the respiratory chain and altering membrane permeability. In our study, we also observed disruption of the redox state, significant increase in intracellular ROS and MDA, and significant decrease in ATP. This demonstrates the biological mechanism by which PCA regulates EMP and PPP, thereby interfering with energy metabolism. The oxidative stress state of the bacteria is not directly related to PCA itself, which is consistent with previous research on the use of PCA as an antioxidant and anti-inflammatory drug in the body [71]. This regulatory mechanism of metabolism can help us discover and screen effective antibacterial candidates.

After learning about the metabolic disorder caused by PCA, we reviewed the synergistic antibacterial mechanism of PCA again. The inhibition of biofilm formation and the change of membrane permeability may be due to this metabolic change, such as the regulation of the formation and dissociation of biofilm by succinic acid and dicarboxylic acid, which is an intermediate product of the citric acid cycle. At the same time, metabolism is also affected by the pentose phosphate pathway. Therefore, whether it is the destruction of EMP by PCA or the blockage of PPP, it will affect the formation of biofilm [90]. Cell membrane permeability is also the same. When the bacterial respiratory chain is damaged by reactive oxygen species, it will cause the oxidation and lipid peroxidation of the cell membrane, thereby reducing the integrity and fluidity of the cell membrane and increasing the permeability of the cell membrane [91]. Therefore, the effect of PCA on metabolism will ultimately be more conducive to the entry of antibiotics into bacteria and play a role, especially the target of carbapenem antibiotics against CR-hvKP itself is on the inner membrane. With the successive collapse of the “natural resistance line” biofilm and cell membrane, the actual efficacy of antibiotics will be greatly improved.

Although we have demonstrated that PCA alters the endogenous oxidative stress of bacteria through metabolic pathways, PCA itself possesses strong reducing properties. Whether this reducing effect will weaken the oxidative stress caused by metabolic changes still needs further investigation. Meanwhile, our study found that under sub-inhibitory concentrations of PCA, the color of the bacterial liquid culture medium deepens with increasing drug concentration. The relationship between this change and metabolism is not clear and requires further exploration.

In conclusion, this study demonstrates that natural plant monomer protocatechuic acid has good antibacterial activity against CR-hvKP and can be used as an antibiotic adjuvant or biofilm inhibitor at low concentrations. It is a promising candidate drug for the development of new antibiotic drugs to combat drug-resistant bacterial pathogens related infections.

## Abbreviations

ATP: Adenosine Triphosphate
BPGM,2,3: Bisphosphoglycerate mutase
c-di-GMP: Cyclic di-GMP
CR-hvKP: Carbapenem-resistant hypervirulent Klebsiella pneumoniae
CRKP: Carbapenem-Resistant Klebsiella pneumoniae
DAPI: 4’,6-diamidino-2-phenylindole
DHFR: Dihydrofolate reductase
DisC3(5): 3,3’-diheptylperoxy carbene-5,5’-dichlorofluorescein
DNA: Deoxyribonucleic Acid
ED: Phosphorylated sugar metabolism pathway
EMP: Glycolytic Pathway
FASR: Fatty Acid Synthesis Reaction
FICI: Fractional Inhibitory Concentration Index
G6PDH: Glucose-6-phosphate dehydrogenase
GAP: Glyceraldehyde-3-phosphate
GC-MS: Gas Chromatography-Mass Spectrometry
HEPES: 4-(2-hydroxyethyl)-1-piperazineethanesulfonic acid
HMGR: Hydroxymethylglutaryl-CoA Reductase
hvKP: Highly Virulent Pneumonia Klebsiella pneumoniae
KCL: Potassium Chloride
LB: Luria-Bertani medium
LC-MS: Liquid Chromatography-Mass Spectrometry
MBC: Minimum Bactericidal Concentration
MDA: Formaldehyde
MIC: Minimum Inhibitory Concentration
NADP(H): Nicotinamide Adenine Dinucleotide Phosphate (reduced form)
NOS: Nitric Oxide Synthase
NR: Nitrate Reductase
NPN: Non-Protein Nitrogen
PCA: Pyrrolidone carboxylic acid
PDAM: 4-phenyl-2,4-diamino-5-methylpyridine
PI: Propyl iodide
PBS: Phosphate Buffered Saline
PPP: Pentose Phosphate Pathway
PYO: Pyrroloquinoline quinone
qRT-PCR: Quantitative Reverse Transcription Polymerase Chain Reaction
R5P: Ribose-5-phosphate
RNA: Ribonucleic Acid
ROS: Reactive Oxygen Species

## Acknowledgements

This study has been made possible thanks to the support from National Key Research and Development Program of China (2023YFD1800100); as well as the joint funding from the National Natural Science Foundation of China (U22A20523).

